# Self-medicating behavior in bumble bees has cascading consequences for pollination and plant reproduction

**DOI:** 10.1101/2025.07.21.665926

**Authors:** Gordon Fitch, Brooke Donzelli, Rebecca E. Irwin, Ken Keefover-Ring, Lynn S. Adler

## Abstract

The sublethal effects of parasites can profoundly influence host traits and propagate to other trophic levels via indirect effects. To date, research on such trait-mediated indirect effects of parasites has focused on non-adaptive changes to host behavior, but adaptive sickness behaviors, such as self-medication, could also indirectly affect community composition and even the evolution of ‘medicinal’ traits in lower trophic levels. Here, we used interactions among the parasite *Crithidia bombi*, the bumble bee host *Bombus impatiens*, and *Monarda fistulosa*, a bee-pollinated plant with multiple chemotypes (genetically determined chemical phenotypes), to experimentally test whether parasite infection influences pollinator foraging, pollination success, and female plant reproduction differentially for medicinal vs. non-medicinal chemotypes.

Compounds from three *Monarda* chemotypes reduced *Crithidia* infection intensity in bees (thymol, carvacrol, and 1,8-cineole; hereafter medicinal chemotypes), while two others did not ((*R*)-(–)-linalool and geraniol; hereafter non-medicinal chemotypes), compared to control sucrose solutions. We found evidence for self-medication in tent foraging choice assays: infected bees preferred medicinal chemotypes while uninfected bees foraged indiscriminately, leading to differences in pollen receipt. *Crithidia* infection had weak but compounding chemotype-specific effects on seed production, germination rate, and offspring chemotype, such that pollination by infected bees resulted in a 57% increase in the proportion of medicinal plants in the F1 generation compared to pollination by uninfected bees. Self-medicating behavior can have differential effects on the reproduction of medicinal vs. non-medicinal plants, suggesting that pollinator parasites may act as agents of selection on the phytochemistry of floral rewards.

**SIGNIFICANCE STATEMENT:** Parasite infection can alter host behavior in multiple ways, including by inducing self-medication. Self-medication has been documented in diverse animal taxa, yet we know very little about the broader ecological or evolutionary consequences of this response to infection. Here, we demonstrate that bumble bees infected with a common parasite show a preference for plant genotypes whose nectar contains antiparasitic compounds, and that this results in differential pollination and reproductive success for medicinal vs. non-medicinal individuals of a chemically polymorphic plant species. Our findings highlight self-medication as a previously understudied mechanism by which parasite infection could initiate cascading effects across trophic levels, and suggest that parasites may indirectly influence the evolution of plant traits via pollinator self-medication behaviors.

## INTRODUCTION

Parasites and pathogens play key roles in regulating host populations (1, 2) and commonly have sublethal effects, including altering host behavior (3, 4). The effects of parasites on hosts can also propagate to other trophic levels either via effects on host density (density-mediated indirect effects, DMIEs) or host traits, especially behavior (trait-mediated indirect effects, TMIEs) (5, 6). In a classic example of a DMIE of pathogens, eradication of the rinderpest virus increased wildebeest populations, which in turn altered fire dynamics, leading to greater tree cover (7). Evidence for TMIEs of pathogens come primarily from aquatic ecosystems; for example, trematode parasites reduce the feeding rate of snail hosts, leading to changes in primary producer biomass and community composition (8, 9). The magnitude of pathogen or parasite-driven TMIEs can rival those of DMIEs (10, 11).

To date, research on indirect effects of parasites has primarily focused on the consequences for antagonistic interactions between hosts and their food sources, but effects on mutualistic interactions may also be significant (12, 13). Moreover, existing research on TMIEs of parasites focuses on the influence of infection on host feeding rate, which is most likely a non-adaptive byproduct of pathogenicity (8, 12). Yet host behavioral responses to infection are diverse and include parasite manipulation, which is adaptive from the perspective of the parasite, and sickness behaviors, which are adaptive for the host (14). The community consequences of these behaviors are largely unexplored.

One key sickness behavior is self-medication, where individuals respond to infection by increasing consumption of a food resource that has a fitness benefit for infected individuals (e.g., by reducing infection), but has neutral or negative effects on the fitness of uninfected individuals (15). Self-medication occurs in a wide range of animal taxa (16–18), but there has been little work investigating whether self-medication-associated shifts in host foraging behavior have measurable effects on the biomass or fitness of medicinal species (but see (19)).

While most well-documented examples of animal self-medication involve herbivory, self-medication may also operate in pollination mutualisms (19, 20). Secondary metabolites found in floral rewards (i.e., nectar and pollen) may have lethal and sublethal negative effects on bees (21, 22), but in some cases they can provide indirect benefits by reducing parasite infection (reviewed in (23, 24)). Bees, in turn, can discern the phytochemical composition of floral rewards, and discriminate among food sources based on phytochemistry (25, 26, but see 27). If bees self-medicate in response to infection by preferentially foraging on plants with antiparasitic properties, parasite infection could lead to cascading effects on plant populations and communities by increasing the fitness of these medicinal plants. However, evidence for self-medication by bees in response to individual infection is rather weak; one field study found that two wild bumble bee species (*Bombus impatiens* and *B. vagans*) infected with the gut parasite *Crithidia bombi* preferred flowers with higher levels of antiparasitic iridoid glycosides (19), while a lab study found that *B. terrestris* workers infected with the same parasite showed a slight preference for sucrose laced with nicotine, a compound with weak antiparasitic effects (20).

Here, we leverage an ecologically important plant-pollinator-parasite system to evaluate whether self-medication by bumble bees (*B. impatiens*) infected with the parasite *Crithidia bombi* influences pollination services, plant reproduction, and the relative frequency of medicinal offspring in a chemically polymorphic plant, *Monarda fistulosa* (wild bergamot or bee balm, Lamiaceae; hereafter, *Monarda*). We first asked whether ecologically relevant doses of the dominant monoterpenes found in the nectar of five *Monarda* chemotypes (genetically determined chemical phenotypes) differentially affect *Crithidia* infection. Finding differences (see Results), we then asked whether foraging behavior of infected vs. uninfected bumble bees differed when presented with a choice between medicinal and non-medicinal *Monarda* chemotypes, and whether this differentially influenced pollination, seed production, germination, and offspring chemotype for medicinal vs. non-medicinal plants (Figure 1). By investigating how self-medication behavior in pollinators affects the reproductive success and trait frequencies of a chemically polymorphic plant, this study reveals a previously overlooked pathway by which parasites can influence pollination dynamics and act as a selective force shaping the evolutionary trajectories of ‘medicinal’ species.

**Figure 1.**
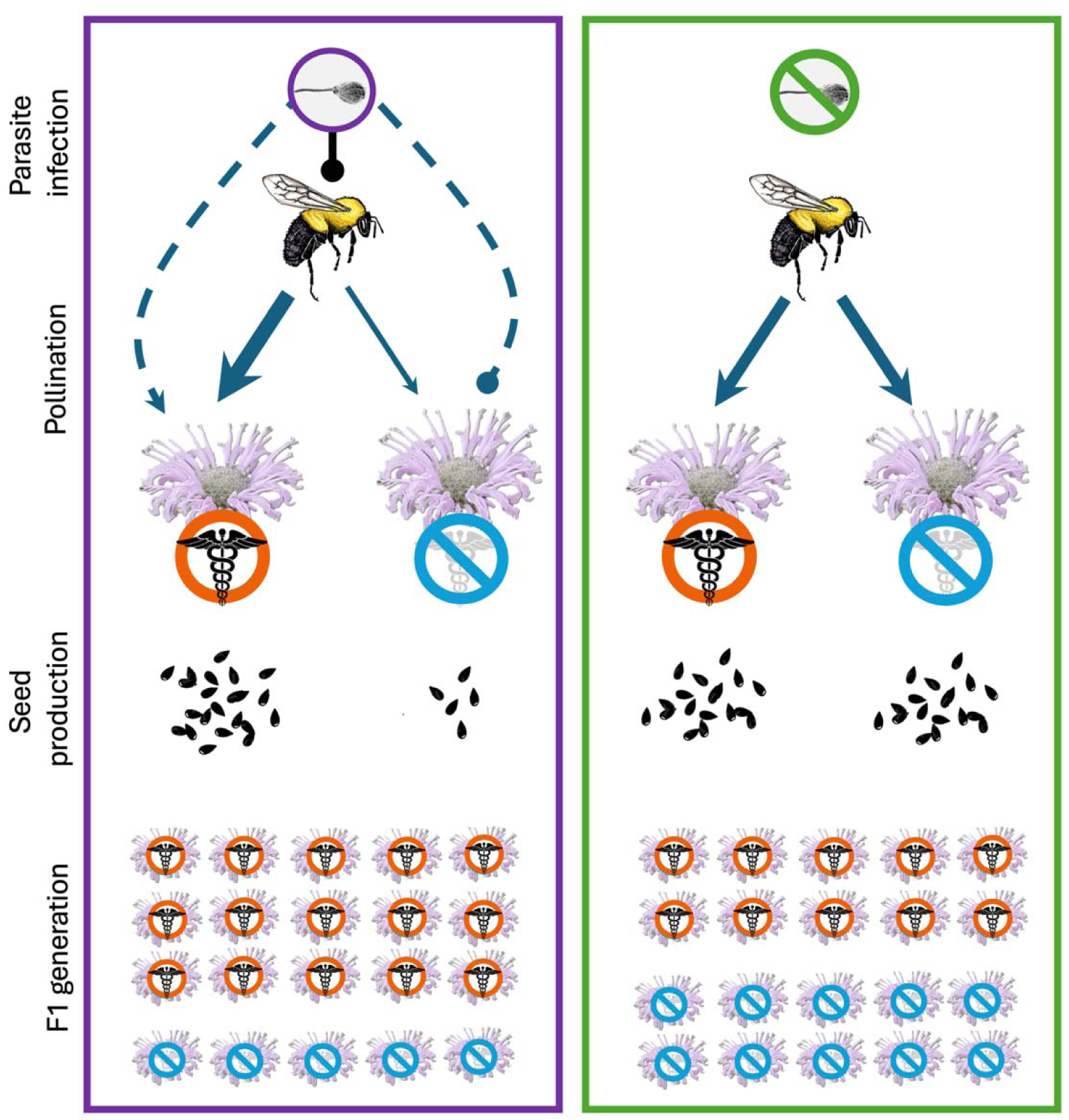
Proposed effects of parasite infection on bumble bee foraging and chemotype relative frequency. Bees infected with *Crithidia* (left) prefer plants whose floral products contain antiparasitic compounds (“medicinal”, red circle and black caduceus) over those that do not (“non-medicinal”, blue circle with gray caduceus), while uninfected bees (right) show no preference. This results in indirect positive effects of *Crithidia* on medicinal plants, which produce more seeds when pollinated by infected bees, and indirect negative effects on non-medicinal plants. This in turn leads to differences in the relative recruitment of medicinal *versus* non-medicinal plants into the F1 generation depending on bee infection status.

## RESULTS

### Do dominant monoterpenes from distinct chemotypes differ in their effect on Crithidia in vivo?

We inoculated lab-reared *B. impatiens* individuals with a known dose of *Crithidia*, then maintained these bees for 7 d on one of six diets [30% sucrose (w/v) with 10 ppm thymol (n = 32), carvacrol (n = 34), (*R*)-(–)-linalool (n = 30; hereafter linalool), geraniol (n = 31), 1,8-cineole (n = 33), or monoterpene-free control (n = 28)], and screened gut contents for *Crithidia* prevalence and infection intensity. *Crithidia* infection differed among diet treatments (^2^ = 35.319, d.f. = 5,158, p < 0.001; *SI Appendix,* Fig. S1) but was not affected by pollen or sucrose consumption rates (^2^ < 0.2, p > 0.7 in both cases). Bees fed thymol, carvacrol or 1,8-cineole diets all showed reduced infection intensity by at least 66% compared to bees fed control sucrose (z > 4.3, p < 0.02 in all cases). Thymol and carvacrol are both phenolic, while 1,8-cineole is not. *Crithidia* infection was not affected by linalool (z = 2.7, p = 0.1) and marginally reduced by geraniol (50% reduction; z = 2.8, p = 0.06) compared to the control sucrose solution. Neither survival nor sucrose or pollen consumption rates differed across diets (survival: ^2^ = 7.45, df = 5,174, p = 0.2; sucrose consumption: F = 0.80, df = 5,184, p = 0.6; pollen consumption: F = 0.87, df = 5,184, p = 0.5).

### Does Crithidia infection mediate foraging behavior on Monarda chemotypes?

To assess foraging preference, in 2022 we conducted trials in which we introduced trios of either *Crithidia*-infected or uninfected *B. impatiens* workers into tents containing a pair of flowering *Monarda* plants of contrasting chemotypes [either one phenolic (i.e., thymol or carvacrol) and one linalool plant (n = 72 trials) or one phenolic and one 1,8-cineole plant (n = 21 trials). Phenolic and 1,8-cineole plants both reduced *Crithidia* infection but are chemically distinct, while linalool plants did not reduce *Crithidia.* This allowed us to assess preference for medicinal vs. non-medicinal chemotypes (phenolic–linalool combination) and between chemically distinct medicinal chemotypes (phenolic–1,8-cineole combination). If bees are self-medicating, infected bees should show a preference for phenolic plants compared to linalool plants, but we expected no differences in preference for uninfected bees, or for any bees offered the choice between phenolic and 1,8-cineole plants, since those chemotypes are both medicinal.

In tents containing one phenolic (i.e., medicinal) and one linalool (i.e., non-medicinal) plant, infected bees were 18% more likely to visit medicinal plants (t = 2.33, d.f. = 312, p = 0.02), while there was no difference in visitation likelihood between chemotypes for uninfected bees (t = 0.46, d.f. = 312, p = 0.6; Table 1, Fig. 2A). Of the plants that were visited, bees spent more time on phenolic plants, regardless of infection (Table 1, Fig. 2B).

**Figure 2.**
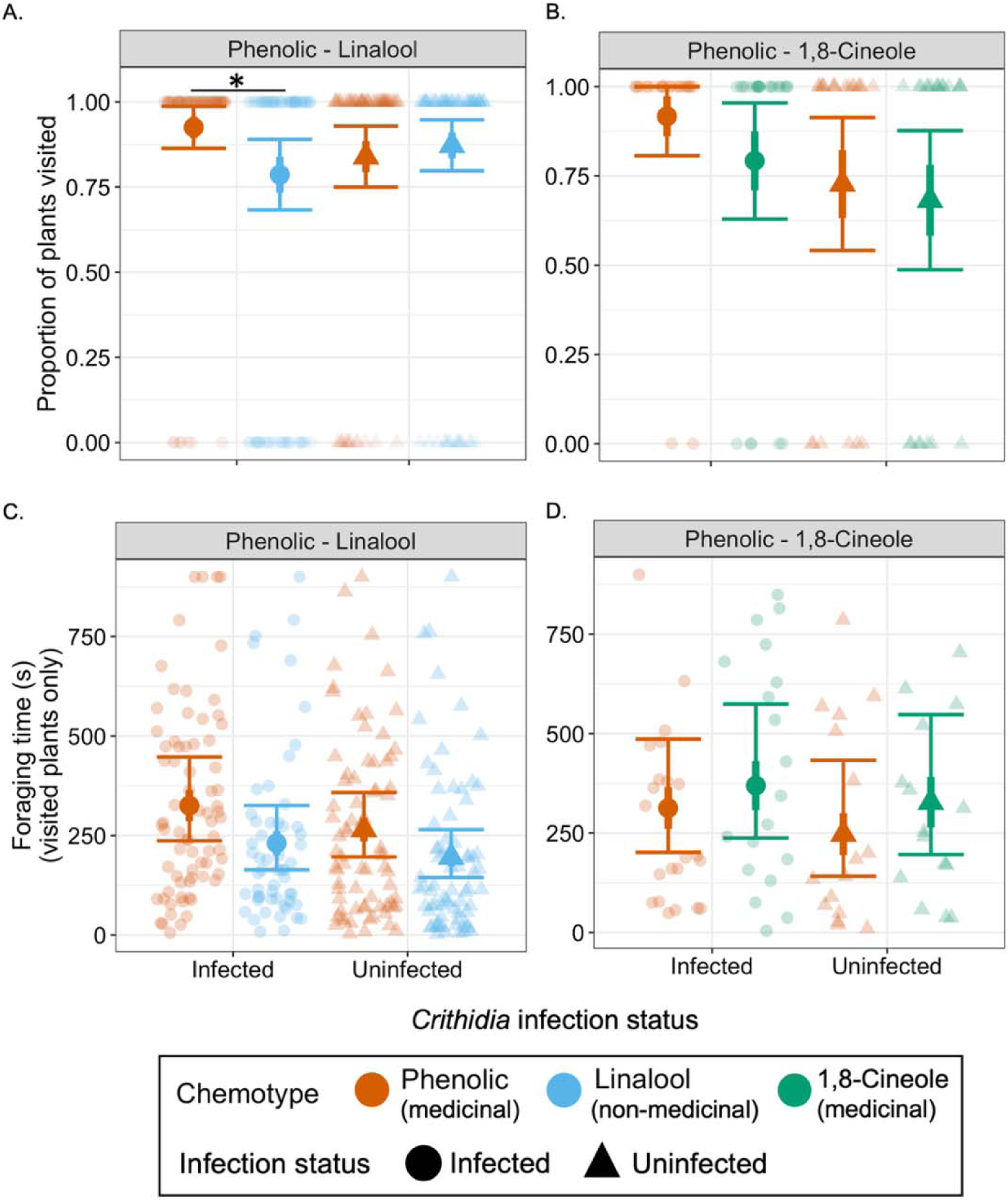
Likelihood of visitation to *Monarda* plants (A, B) and time spent foraging (C, D) by infected and uninfected bees in phenolic-linalool (i.e., medicinal vs. non-medicinal) tents (A, C) and phenolic-1,8-cineole (i.e., phenolic medicinal vs. non-phenolic medicinal) tents (B, D), two-plant experiment. Faint points represent observations; large points represent means, with thick bars indicating ± 1SE and whiskers indicating 95% CIs. *p < 0.05. In C, foraging time on phenolic plants was significantly higher than on linalool plants regardless of bee infection (p = 0.02). If bees self-medicate, then we expect to see that infection status alters behavior in medicinal vs. non-medicinal comparisons (A, C) but not medicinal vs. medicinal (B, D).

**Table 1.**
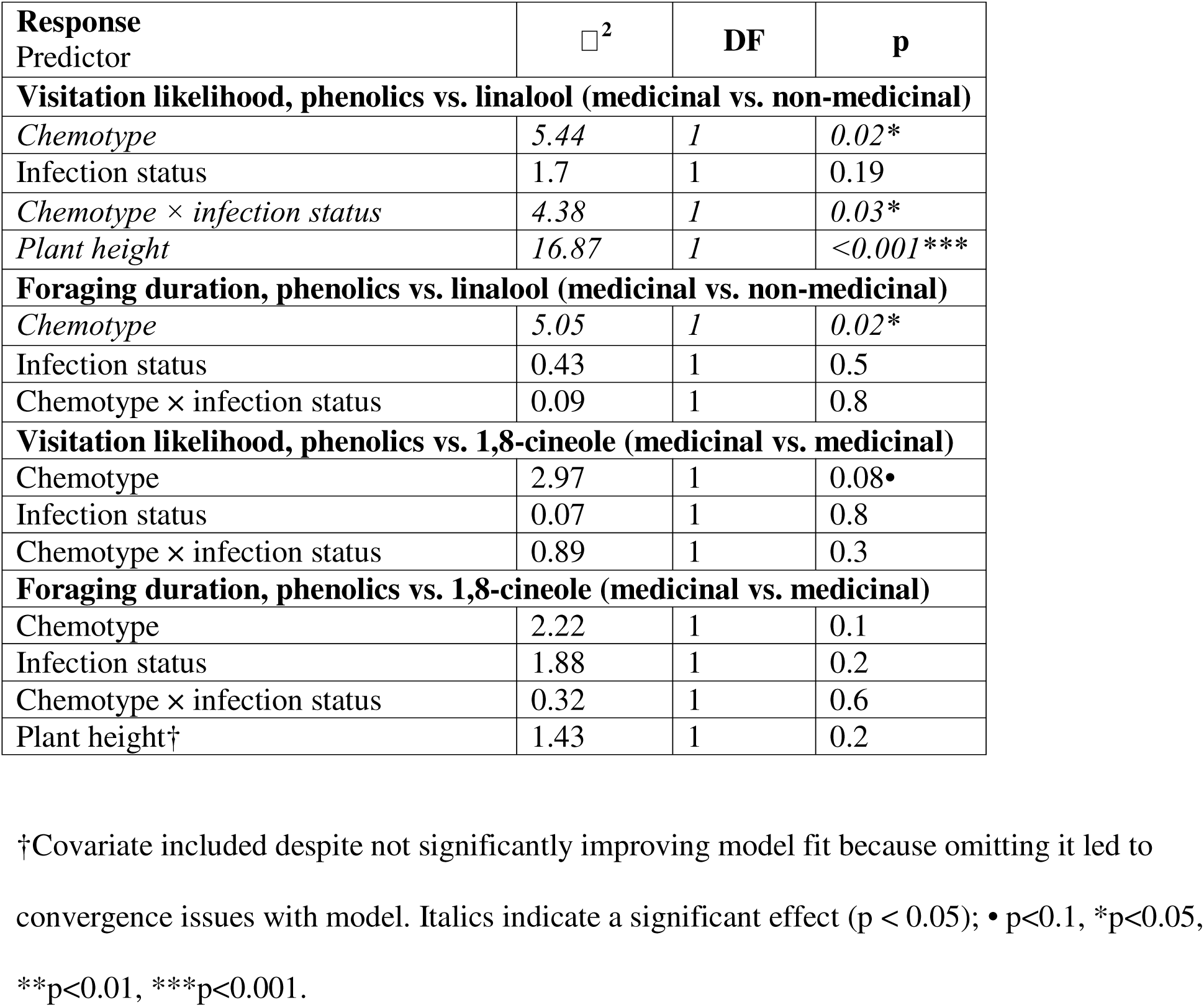
Interactive effects of chemotype and infection status on bumble bee foraging behavior, from the two-plant tent experiment (conducted 2022). Consumption of phenolic compounds and 1,8-cineole reduce *Crithidia* infection in bumblebees but have different chemical structures and scents; linalool does not reduce *Crithidia*. We therefore predicted that if bees self-medicate, infection status would affect how bees respond to medicinal vs. non-medicinal chemotypes, but would not affect bee behavior when choosing between two medicinal chemotypes.

In tents containing a 1,8-cineole plant and a phenolic plant (i.e., choice between two medicinal plants), there was no effect of any predictor on the likelihood of visitation (Table 1). Similarly, we detected no differences in time spent foraging on plants based on chemotype or infection (p > 0.1 in all cases; Table 1, Fig. 2D).

### Does infection-mediated foraging behavior differentially affect pollination and plant female reproduction across chemotypes?

#### Pollination

After foraging trials concluded, we assessed stigma pollen load (number of *Monarda* pollen grains on stigma) on a subset of flowers (n = 159). Outside of trials, flowers were bagged to preclude visitation. There was no effect of any predictor on the likelihood of a stigma receiving pollen (p > 0.5 for all predictors, Fig. 3A). However, for stigmas that did receive pollen, there was a significant chemotype × infection status interaction, though the main effects of chemotype and infection status on pollen load were not significant (Table 2). The significant chemotype × infection status interaction was driven by a substantial difference in pollen load for plants visited by uninfected bees, with non-medicinal linalool plants receiving >4x more pollen than medicinal phenolic plants (3.4±1.6 vs. 0.8±0.4 pollen grains per stigma; z = 2.59, d.f. = 46, p = 0.009); there was no parallel difference for flowers visited by infected bees (2.2±1.0 vs. 3.0±1.3 pollen grains per stigma; z = -0.62, d.f. = 46, p = 0.5; Fig. 3B).

**Figure 3.**
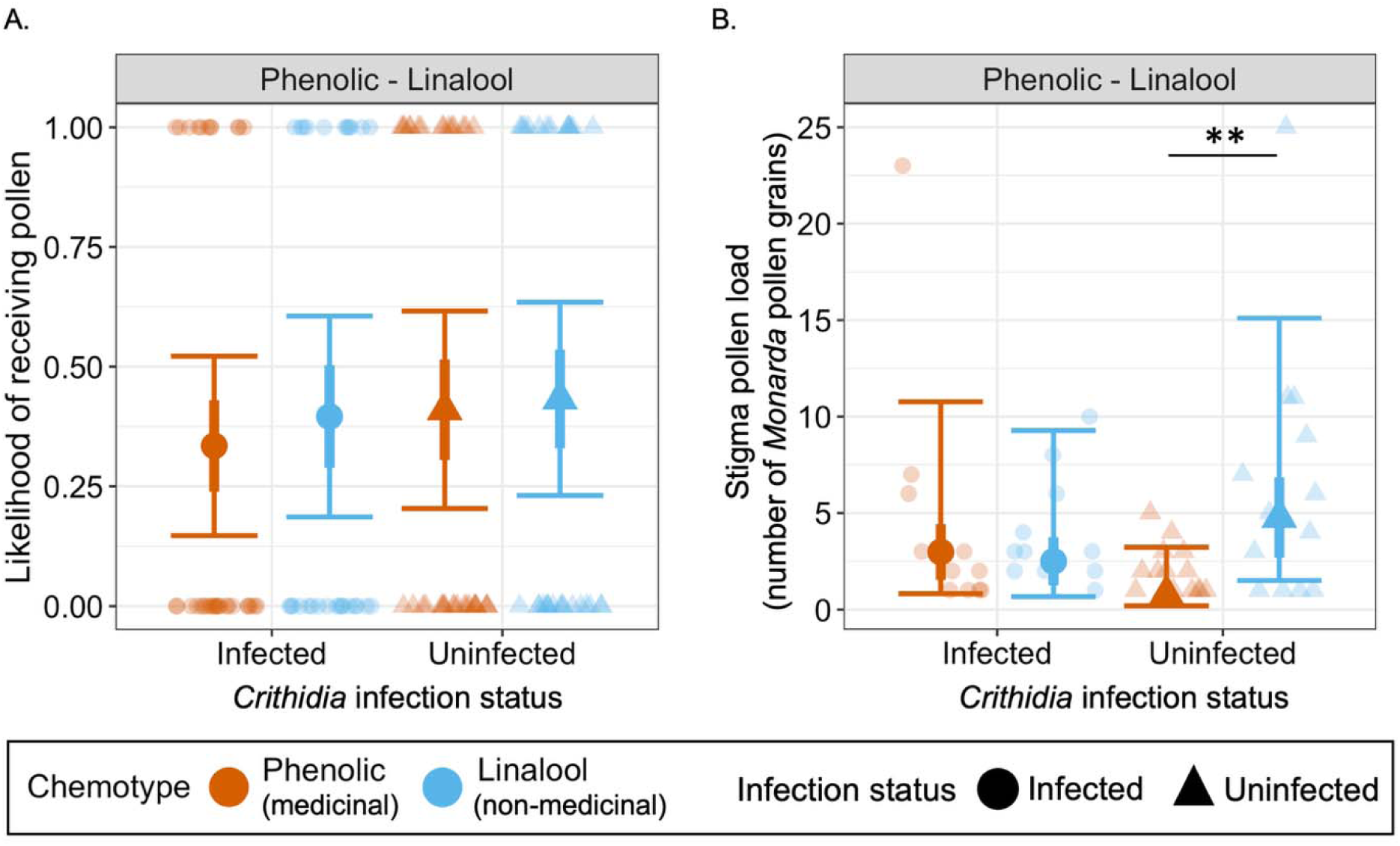
Likelihood of a stigma receiving pollen (A) and number of *Monarda* pollen grains on stigmas that received pollen (B) based on infection status of pollinating bees, two-plant experiment. Faint points represent observations; large points represent means, with thick bars indicating ± 1SE and whiskers indicating 95% CIs. **p < 0.005.

**Table 2.** Interactive effects of chemotype and infection status on stigma pollen load (*Monarda* pollen grains per stigma) from two-plant tent experiment (conducted 2022).

| **Response**<br>Predictor | $\chi^2$ | **DF** | **p** |
| --- | --- | --- | --- |
| **Likelihood of having $\geq 1$ pollen grain, phenolics vs. linalool** |  |  |  |
| Chemotype | 0.60 | 1 | 0.4 |
| Infection status | 0.75 | 1 | 0.4 |
| Chemotype × infection status | 1.92 | 1 | 0.2 |
| **Stigma pollen load, phenolics vs. linalool** |  |  |  |
| Chemotype | 1.65 | 1 | 0.2 |
| Infection status | 1.00 | 1 | 0.3 |
| *Chemotype × infection status* | *5.40* | *1* | *0.02\** |

Italics indicate a significant effect ( $p < 0.05$ ); • $p < 0.1$ , \* $p < 0.05$ , \*\* $p < 0.01$ .

### Seed production, germination, and offspring chemotype

To assess how differences in pollinator visitation and pollination translated to seed production, in 2023 we conducted a similar experiment but increased the number of plants per tent to 6 and increased the time flowers were exposed to bees >30-fold. We focused on plant reproduction this year and did not conduct behavioral observations or measure pollen receipt. From these tents, we collected 1062 inflorescences to measure seed production (X = 14.75 inflorescences plant^-1^, range = 2-40) and retrieved 79012 seeds (X = 74.4 seeds inflorescence^-1^, range = 0-354). Seeds per inflorescence was 15% lower for plants pollinated by infected vs. uninfected bees and was positively correlated with inflorescence diameter; there was no effect of chemotype or chemotype X infection status (Table 3; Fig. 4A). For total seeds per plant, by contrast, there was no main effect of either chemotype or infection status, but the chemotype X infection status interaction was marginally significant (^2^ = 2.90, df = 1,64, p = 0.08); this interaction was consistent with foraging data from the 2022 experiment, with thymol plants producing more seeds than linalool plants in tents with infected bees (estimated marginal means: 2192 ± 464 vs. 1745 ± 416), and the opposite in tents with uninfected bees (2055 ± 441 vs. 2444 ± 504) (Table 3; Fig. 4B). Per-seed mass was not influenced by any predictor (Table 3).

**Figure 4.**
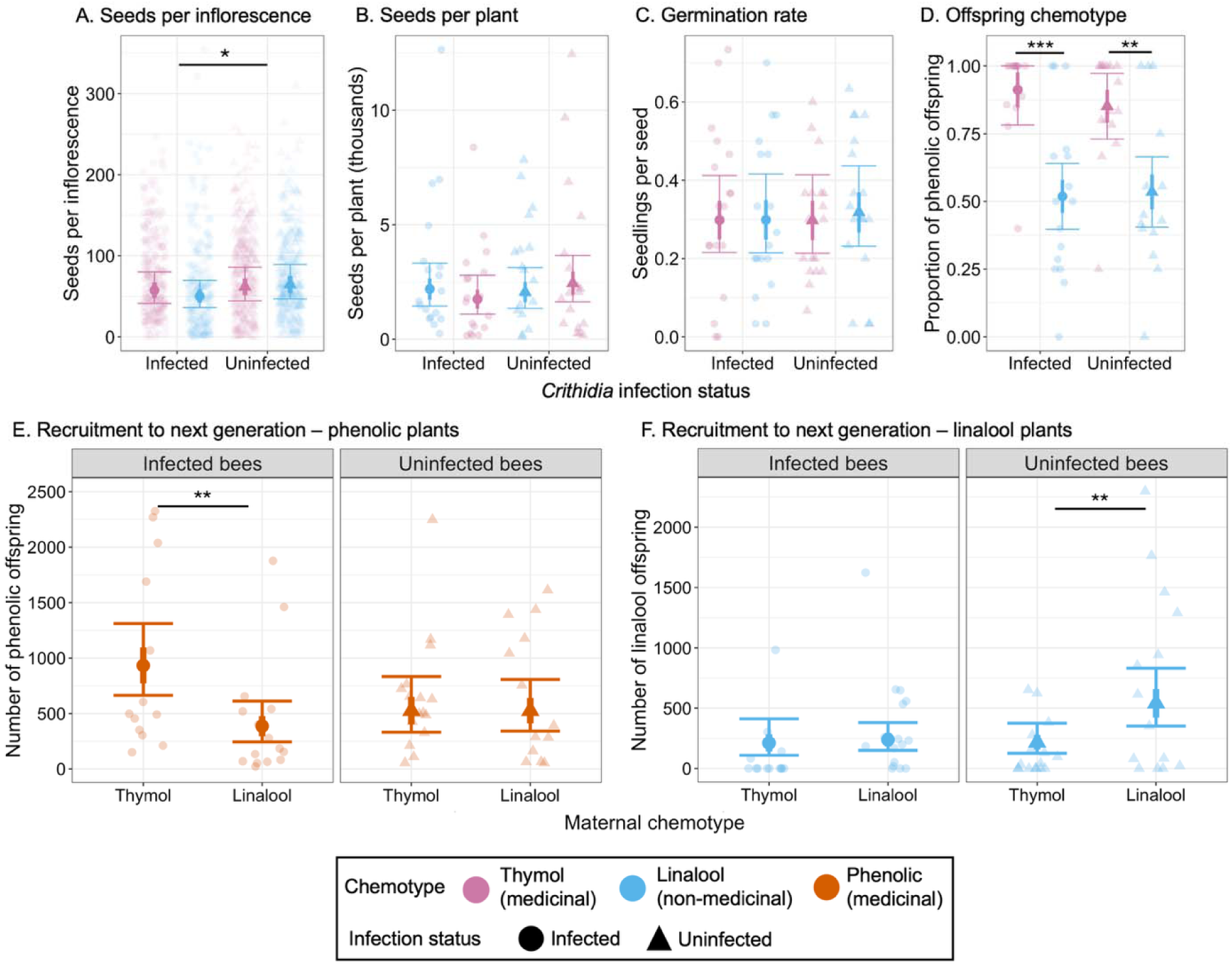
Reproductive outcomes from the six-plant tent experiment comparing thymol and linalool plants pollinated by infected and uninfected bees: A) per-inflorescence seed production, B) per-plant seed production, C) germination rate, D) proportion of phenolic offspring, E) recruitment of phenolic plants to next generation, and F) recruitment of linalool plants to next generation. Panels E and F represent the product of values from panel B X panel C X panel D. Faint points represent observations; large points represent means, with thick bars indicating ± 1SE and whiskers indicating 95% CIs. *p < 0.05; **p < 0.005, ***p < 0.001.

**Table 3.** Interactive effects of chemotype and infection status on *Monarda* seed production, germination, and offspring chemotype, six-plant experiment (conducted 2023). All trials compared thymol (medicinal) plants to linalool (non-medicinal) plants.

| Response Predictor | $\eta^2$ | DF | p |
| --- | --- | --- | --- |
| **Seeds per inflorescence** |  |  |  |
| Chemotype | 0.48 | 1, 1041 | 0.5 |
| *Infection status* | 4.52 | 1, 1041 | 0.03* |
| Chemotype × infection status | 1.19 | 1, 1041 | 0.28 |
| *Inflorescence diameter* | 248.21 | 1, 1041 | <0.001*** |
| *Observer* | 65.57 | 6, 1041 | <0.001*** |
| **Seeds per plant** |  |  |  |
| Chemotype | 0.73 | 1, 64 | 0.4 |
| Infection status | 2.19 | 1, 64 | 0.1 |
| Chemotype × infection status | 2.90 | 1, 64 | 0.09• |
| *Mean inflorescence diameter* | 18.92 | 1, 64 | <0.001*** |
| **log(Per-seed mass)** |  |  |  |
| Chemotype | 0.06 | 1, 1012 | 0.8 |
| Infection status | 0.11 | 1, 1012 | 0.7 |
| Chemotype × infection status | 3.30 | 1, 1012 | 0.07• |
| *Mean inflorescence diameter* | 12.43 | 1, 1012 | <0.001*** |
| *Observer* | 15.85 | 6, 1012 | 0.01* |
| **Germination rate** |  |  |  |
| Chemotype | 0.07 | 1, 64 | 0.8 |
| Infection status | 0.03 | 1, 64 | 0.9 |
| Chemotype × infection status | 0.06 | 1, 64 | 0.8 |
| *Plant height* | 6.19 | 1, 64 | 0.01* |
| **Proportion of phenolic offspring** |  |  |  |
| *Chemotype* | 59.28 | 1, 54 | <0.001*** |
| Infection status | 0.09 | 1, 54 | 0.8 |
| Chemotype × infection status | 0.30 | 1, 54 | 0.6 |
| **Estimated number of offspring, by chemotype** |  |  |  |
| Maternal chemotype | 1.98 | 1, 108 | 0.2 |
| *Infection status* | 9.53 | 1, 108 | 0.002** |
| *Maternal chemotype* × | 10.10 | 1, 108 | 0.001** |
| *infection status* |  |  |  |
| Offspring chemotype | 1.98 | 1, 108 | 0.2 |
| *Maternal chemotype × offspring chemotype* | *39.03* | *1, 108* | *<0.001\*\*\** |
| *Mean inflorescence diameter* | *16.44* | *1, 108* | *<0.001\*\*\** |
| *Plant height* | *15.76* | *1, 108* | *<0.001\*\*\** |
| **Estimated number of phenolic offspring** |  |  |  |
| *Maternal chemotype* | *9.84* | *1, 52* | *0.002\*\** |
| *Infection status* | *1.27* | *1, 52* | *0.3* |
| *Chemotype × infection status* | *5.73* | *1, 52* | *0.02\** |
| *Plant height* | *14.03* | *1, 52* | *<0.001\*\*\** |
| *Mean inflorescence diameter* | *9.47* | *1, 52* | *0.002\*\** |
| **Estimated number of linalool offspring** |  |  |  |
| *Maternal chemotype* | *0.09* | *1, 50* | *0.8* |
| *Infection status* | *36.24* | *1, 50* | *<0.001\*\*\** |
| *Chemotype × infection status* | *6.64* | *1, 50* | *0.0099\*\** |
| *Plant height* | *12.65* | *1, 50* | *<0.001\*\*\** |
| *Zero inflation ~ Maternal chemotype* | *2.58* | *1, 50* | *0.0098\*\** |

Italics indicate a significant effect (p < 0.05); • p<0.1, \*p<0.05, \*\*p<0.01, \*\*\*p<0.001.

Bee infection status strongly impacted the chemotype composition of the F1 generation. The estimated number of phenolic (i.e., medicinal; thymol or carvacrol) offspring per plant was 55% higher for thymol vs. linalool mothers and, while there was no main effect of infection, there was a significant chemotype X infection status (Table 3, Fig. 4E). In contrast, linalool (i.e., non-medicinal) offspring per plant did not differ between chemotypes, but was 52% higher in plants visited by uninfected bees (Table 3, Fig. 4F). Post-hoc tests indicated that production of phenolic offspring was higher for thymol plants than linalool plants when pollinated by infected bees (z = 3.14, p = 0.002), but similar in tents with uninfected bees (z = 0.002, p > 0.9; Fig. 4E), while production of linalool offspring was equal between thymol and linalool plants in infected tents (z = -0.31, p = 0.8) but higher for linalool plants in uninfected tents (z = 2.60, p = 0.009; Fig. 4F). These differences mean that pollination by infected bees resulted in thymol plants producing an estimated 77% more phenolic offspring per plant compared to pollination by uninfected bees (933±162 vs. 526±124; Fig. 4E), while linalool plants produced 56% fewer linalool offspring when pollinated by infected bees (239±56 vs. 541±119; Fig. 4F). This outcome reflects subtle chemotype X infection status signatures on both germination rate and offspring chemotype that, while not approaching statistical significance on their own, reinforce the overall pattern of medicinal plants having higher fitness when pollinated by *Crithidia-*infected than uninfected bees and *vice versa* (Table 3, Fig. 4C-D, *SI Appendix*, S.2.2).

## DISCUSSION

We found that infection with the gut parasite *Crithidia* altered bumble bee foraging preferences, with infected bees more likely to visit *Monarda* plants that produced antiparasitic compounds (i.e., medicinal plants) than non-medicinal plants. This translated into differences in pollen deposition, with medicinal plants receiving more pollen than non-medicinal plants in tents with infected bees, and *vice versa* for tents with uninfected bees. Small individual effects of differential pollination on seed production, germination, and the chemotype of resulting offspring meant that recruitment of medicinal-chemotype plants into the next generation was significantly higher when plants were pollinated by infected bees than uninfected bees, while recruitment of non-medicinal plants was increased in tents pollinated by uninfected bees. This is consistent with a growing body of evidence that parasite infection can lead to TMIEs on lower trophic levels (5, 6, 8, 12), and demonstrates that these cascading effects can ultimately drive evolution in a resource species.

Previous studies on TMIEs of parasites have focused on the impact of altered host feeding rates (8, 10, 11), but here we show that changes in feeding preference can have similar cascading effects and drive evolution of chemical phenotypes. According to our results, a *Monarda* population with a 1:1 ratio of medicinal to non-medicinal chemotypes would, if pollinated exclusively by infected bees, produce 1145±178 medicinal plants compared to 625±107 non-medicinal plants per medicinal/non-medicinal pair, giving a ratio of 1.8:1 in the F1 generation. If, on the other hand, the plants were pollinated exclusively by uninfected bees, they would produce 744±138 medicinal plants vs. 1067±166 non-medicinal plants, for a medicinal-to-non-medicinal ratio of 0.7:1. Thus, the prevalence of *Crithidia* infection in bumble bees could determine whether the relative abundance of medicinal-medicinal plants would increase, decrease, or stay constant through time.

In addition to the self-medication-mediated indirect effects (IEs) of parasite infection, we also found evidence for a feeding efficiency-mediated IE (10, 30, 31). *Monarda* plants produced fewer seeds per inflorescence when visited by infected bees, regardless of chemotype. This is consistent with research showing that *Crithidia* infection impairs foraging efficiency in bumble bees (32, 33), but to date there has been scant evidence that such effects can depress pollination services and plant reproduction (but see ref. 13). However, this inflorescence-level effect did not scale up to influence plant-level seed production, suggesting that *Monarda* may compensate for reduced pollination by producing more inflorescences. Similar compensatory responses to low pollination have been observed in other species, but likely depend on resource availability and plant condition (34, 35).

It is not yet clear whether the TMIEs of *Crithidia* we observed in tent experiments are strong enough to influence wild *Monarda* populations. On the one hand, among plants in our 2022 study of pollination dependence, open-pollinated medicinal (phenolic) inflorescences received significantly more pollen than open-pollinated non-medicinal (linalool) inflorescences (*SI Appendix*, S2.2). *Crithidia* prevalence can be well over 50% in bumble bees in our study region during the *Monarda* bloom period (13, 36); bumble bees were the most common visitors to open-pollinated inflorescences and were observed transporting *Monarda* pollen (G. Fitch, personal observation). High levels of *Crithidia* infection in wild bumble bees therefore may have contributed to more pollination in plants producing antiparasitic phenolic compounds (though this did not translate into more seeds produced per inflorescence; *SI Appendix*, S2.2). On the other hand, the increased complexity of natural plant-pollinator-parasite communities may reduce the impact of TMIEs of pollinator parasites. Access to diverse floral resources with antiparasitic properties may dampen the impact of self-medicating behavior on any single plant species. Moreover, in diverse pollinator communities, pollinator species will likely differ in susceptibility to parasite infection (37, 38), and antiparasitic effects of floral products may be host species-specific (39), all of which could attenuate behavior-mediated IEs of parasites.

Further research in field conditions and more species-rich controlled experiments are needed to determine the role of pollinator parasites in shaping the species and phytochemical compositions of flowering plant communities.

The differences in foraging behavior we observed between infected and uninfected bees are consistent with self-medication. However, because we did not investigate bee fitness in this study, we cannot definitively say that bees were self-medicating (15). Future studies should quantify the fitness impact of parasitism-induced changes to foraging behavior, as well as the fitness effects for uninfected bees of diets including compounds with antiparasitic effects. Given the challenges in assessing the fitness effects of individual foraging choices in eusocial species such as bumble bees (18), future studies could assess proxies for fitness [e.g., individual survival (20)] or employ microcolonies to increase tractability (28).

### Conclusion

This study demonstrates parasite-induced changes to foraging behavior of bumble bees consistent with self-medication, and that these changes can differentially influence pollination and reproduction of medicinal vs. non-medicinal genotypes of *Monarda*. Our findings highlight self-medication as a previously understudied mechanism by which parasite infection could initiate TMIEs across trophic levels, and suggest that TMIEs of parasites may influence not only primary producer community composition, but also the evolution of plant traits relevant to self-medication behaviors.

## METHODS

### Study system

*Monarda fistulosa* L. is a perennial, bumble bee-attracting wildflower that produces monoterpenes in glandular trichomes, mainly on leaves and floral structures. *Monarda* populations comprise multiple genotypes that differ in their dominant monoterpene (i.e., chemotypes); chemotypes include thymol and carvacrol (isomeric phenolic compounds), (*R*)-(–)-linalool, geraniol, and 1,8-cineole (29) (see *SI Appendix*, S1.1 for additional details).

Morphologically, flowers of the different chemotypes are indistinguishable (*SI Appendix*, Table S2). Flowers are protandrous, limiting autogamy (40), and plants can produce multiple flowering inflorescences in a single season.

*Bombus impatiens* is the most common bumble bee species across much of eastern North America (41). Bumble bees, including *B. impatiens*, are among the most frequent visitors to *Monarda* (40, 42). *Crithidia bombi* is a common and widespread, horizontally-transmitted gut parasite of bumble bees [e.g., prevalence of up to 80% at some sites in the northeastern USA (36)]. *Crithidia* infection has negative consequences for bumble bee queen fitness, worker survival, and colony performance, particularly in resource-stressed bees (43, 44), and has been implicated in the decline of some bumble bee populations (45).

### Do dominant terpenes from distinct chemotypes differ in their effect on *Crithidia in vivo*?

#### Diet solutions

We assessed thymol, carvacrol, (*R*)-(–)-linalool [the dominant enantiomer in linalool-type *Monarda* plants (29)], geraniol, and 1,8-cineole (Sigma Aldrich, Saint Louis, MO USA *SI Appendix*, Table S3). Each was incorporated at 10 ppm into a 30% sucrose solution (*SI Appendix,* S1.2.1). Ten ppm lies within the range of concentrations recorded for thymol in the nectar of thyme (*Thymus vulgaris*; Lamiaceae) (46, 47), a plant with similar chemotypes to *Monarda* that likely arise via the same biosynthetic pathway (29, 48). After this study was completed, we analyzed single *Monarda* nectar samples from thymol and linalool plants, and found that nectar monoterpene concentration is comparable to the levels tested here (6.1 ppm for thymol and 10.4 ppm for linalool; *SI Appendix*, S1.6). Therefore, we are confident that our findings regarding the effects of each chemotype are ecologically relevant.

#### *Diet assays and* Crithidia *screening*

We conducted six trials between 20 Oct and 10 Nov 2021. In each trial, 36 bees were inoculated with *Crithidia* and fed one of six diets (2 bees per colony per diet for each trial). Experimental bees came from three *Crithidia*-free commercial colonies (Koppert Biological, Lansing, MI, USA), which were maintained in darkness at 28 °C on a diet of 30% sucrose and wildflower pollen (lot #81001, CC Pollen Company, Phoenix, AZ, USA). We infected experimental bees by feeding them an inoculum containing 9000 *Crithidia* cells in 25% sucrose solution, following standard protocols (28, 47) (*SI Appendix*, S1.2.4). Inoculated bees were randomly assigned to one of six diet treatments [30% sucrose (w/v), plain (control) or with 10 ppm thymol, carvacrol, linalool, geraniol, or 1,8-cineole] and placed into individual 473 ml clear plastic feeding cups (Dart Containers, Mason, MI, USA), supplied with 10 ml of the assigned diet treatment in a nectar feeder (made from a 35 mm petri dish with a dental cotton wick) and a 0.5 g ball of wildflower pollen. All bees were maintained in darkness at 28 °C for the duration of the experiment. Over the first 48 h, we measured consumption of sucrose solution and pollen, to see whether this varied among diet treatments and influenced infection (*SI Appendix*, S1.2.2). Nectar feeders and pollen were replaced every 2 d, at which time we also removed any dead bees, recording day of removal and marginal cell length on the right forewing for each [a proxy for body size (49)]. After 7 d, at which point *Crithidia* infection has reached a representative level (50), we sacrificed bees and screened gut contents for *Crithidia* using established protocols (47) (*SI Appendix*, S1.2.3). Because body size can influence *Crithidia* infection intensity (51), we measured marginal cell length of each bee as a proxy for size.

#### Data Analysis

All analyses were conducted with R version 4.3.3 (52). We compared *Crithidia* cell count (cells/0.02 µL) across treatments with a negative binomial generalized linear mixed model (GLMM) (see *SI Appendix,* Table S4 for full formulation for all models). We used a post-hoc pairwise Tukey’s test with the package ‘emmeans’ (53) to assess differences in *Crithidia* cell count across treatments. We compared evaporation-corrected sucrose consumption across treatments using a linear mixed-effects model (LMM). We tested whether diet treatment affected mortality using a Cox proportional hazards model, implemented using ‘coxph()’ from the ‘survival’ package (54), with marginal cell length and colony as covariates. For all models, we used stepwise model simplification with likelihood-ratio tests to find the most parsimonious model (always retaining predictors of interest), and for the most parsimonious model, we checked that data conformed to model assumptions using ‘DHARMà (55).

### Does *Crithidia* infection mediate foraging behavior on *Monarda* chemotypes?

#### Site preparation

Trials were conducted at the University of Massachusetts Crop and Animal Research and Education Farm (South Deerfield, MA, USA: 42.475149, -72.582279). Plants >1 year old were transplanted in July 2021 or June 2022 (see *SI Appendix*, S1.1 for details on plant sources). In June 2022, plants were assigned to one of 36 1.8m x 1.8m plots, each of which contained a central linalool or 1,8-cineole plant surrounded by four phenolic plants. To confirm the chemotype of all plants, we conducted gas chromatography on a single leaf of each plant (29). Plants were watered as needed, and plots were mulched and periodically weeded.

#### Foraging behavior

In 2022, we observed foraging activity of trios of infected or uninfected bees presented with a choice of two *Monarda* plants of different chemotypes in mesh tents (0.68 m^3^ or 0.75 m^3^, depending on plant size) erected over the plants. We used *B. impatiens* workers from three commercial colonies (Koppert Biological: Lansing, MI USA). To control for the effects of between-colony variation, we evenly divided each colony into infected and uninfected subcolonies that were used to supply workers for trials (*SI Appendix*, S1.3.1). In 72 trials, paired plants included one non-medicinal (linalool) and one medicinal (phenolic) plant. In 21 additional trials, paired plants included one 1,8-cineole and one phenolic plant; 1,8-cineole has comparable antiparasitic effects to thymol and carvacrol (see Results) but different chemical structure and scent. We included the 1,8-cineole-phenolic comparison to check whether infection-mediated differences in behavior were driven by differential antiparasitic activity or simply chemical dissimilarity.

Observations took place between 0900-1600 h from 07 Jul to 04 Aug 2022. Trios of color-marked bees were introduced to a tent and tracked by two observers. Once at least one bee had foraged on both plants, the observers began a 15 min observation period of all three bees.

During the observation period, all foraging visits (defined as uninterrupted foraging on a single plant) were noted separately for each bee. For each visit, the observer tallied the number of inflorescences probed (hereafter ‘probes’) and timed per-visit duration using a stopwatch (*SI Appendix*, S1.3.3).

Bees were left to forage for 1 h from first foraging, after which all bees were removed from the tent and returned to their cup, provisioned with 30% sucrose solution *ad libitum*, returned to the lab, and held at 27 °C overnight. The following morning, we screened gut contents of each bee for *Crithidia* infection intensity.

### Does infection-mediated foraging behavior differentially affect pollination and plant female reproduction across chemotypes?

#### Pollination

In the 2022 experiment, we assessed stigma pollen load on a subset of inflorescences that were only accessible to pollinators during a single foraging trial. These inflorescences were enclosed in a 13 cm x 17 cm mesh bag (Uline, Pleasant Prairie, WI, USA) prior to flowering.

Inflorescences were unbagged immediately before a foraging trial began. After the trial concluded, we collected one randomly selected receptive stigma from each unbagged inflorescence before replacing the bag. The harvested stigma was transferred to a microcentrifuge tube and kept on ice. Upon returning to the lab, we mounted each stigma on a microscope slide with basic fuchsin gel (56) and counted *Monarda* pollen grains under a compound microscope at 100x.

To assess the degree of pollinator dependence for pollination and seed set, we also compared pollination and seed production from these inflorescences to that seen in bagged inflorescences (no pollination) and to fully accessible (unbagged) inflorescences on the same plants (*SI Appendix*, S1.7 & S2.2).

#### Seed production

In 2022 trials, pollination was insufficient to influence seed production (*SI Appendix,* Fig. S2, Table S1). Therefore, in 2023 we created 12 modified plots with 3 medicinal (thymol) and 3 non-medicinal (linalool) plants each. Plots were paired, and for each pair of plots, 6 plants from the 2022 study were dug up, split in half, and transplanted back into the ground, such that the plants in one tent were genetically identical to the corresponding plants in its paired tent. After transplanting, we erected 0.75 m^3^ mesh tents over each plot; these remained in place for the duration of flowering. In each pair of plots, one tent was randomly assigned to infected bees and the other to uninfected bees.

Once flowering began, we introduced groups of 5 infected or uninfected worker bees to each tent for 3 h each day, 5 days per week, for the duration of the flowering period. We confirmed that bees were foraging but did not record behavior. After the 3 h foraging period, we removed all bees from the tent; new bees were used each day to limit the effects of learning.

After the flowering period concluded, we enclosed all inflorescences in individual mesh bags to deter seed predators, up to 50 inflorescences per plant. Once inflorescences matured, we removed them from the plant, leaving them in the mesh bag to contain any dislodged seeds. We subsequently measured the diameter of the mature inflorescence to include as a covariate, counted the number of seeds, and weighed the bulk seeds by inflorescence to the nearest 0.01 g. We then calculated weight per seed by dividing bulk weight by the number of seeds.

#### Seed viability and offspring chemotype

Using seeds from the 2023 experiment, we assessed germination rate and the chemotype of germinated offspring for each maternal plant by sowing 30 randomly selected seeds per plant in 200-cell germination trays (Jiangsu Grow-Green, Jiangsu, China) filled with York University Greenhouse potting soil mix (*SI Appendix*, Table S5). Trays were maintained under greenhouse conditions with ambient light and regular watering until plants reached the seedling stage, at which point we transplanted seedlings to 14 cm square pots (JMCS55-1, McConkey, Saint-Léon-de-Standon, QC, Canada). Each plant was chemotyped by scent (linalool or phenolic) by one experienced observer (GF), and confirmed by a second observer (Brandi Zenchyzen), once they had at least 10 pairs of leaves. We validated chemotype classifications by analyzing leaves from 10% of plants via GC; all scent-based classifications matched GC classifications.

#### Data analysis

For analysis of the 2022 tent experiment, we grouped thymol and carvacrol plants together as phenolic plants, given their similar chemical structure and effects on *Crithidia*. An alternative analysis considering thymol and carvacrol plants separately yielded qualitatively similar results (*SI Appendix,* Table S6).

To analyze how bee visitation was affected by infection status and plant chemotype, we included only bees that foraged during the observation period. We used a two-step hurdle model to account for excess zeroes in the dataset. First, we modeled whether a bee visited a plant (binary response) using a binomial generalized mixed-effects model (GLMM). Then, for plants that received at least one visit, we modeled three aspects of visitation per bee (total foraging time summed across visits, number of visits, and number of probes), using three GLMMs, each with a zero-truncated negative binomial error distribution. Because patterns were qualitatively similar for foraging time, number of visits, and number of probes (*SI Appendix*, Table S7), we discuss only foraging time. All models were implemented using the glmmTMB package (see *SI Appendix*, Table S4 for model formulations). Stigma pollen loads, like visitation data, were strongly zero-inflated, so we again used hurdle models to analyze whether stigma pollen load was influenced by chemotype, bee infection status and their interaction. We analyzed only phenolic-linalool tents due to low sample size for phenolic-1,8-cineole tents (N = 16 1,8-cineole stigmas, of which 4 received pollen).

We assessed female reproductive success in the 2023 experiment via 1) seeds per inflorescence, 2) seeds per plant and 3) per-seed mass, asking whether female reproductive success differed by plant chemotype, bee infection status, and chemotype × infection interaction. We calculated seeds per plant by taking the mean number of seeds per inflorescence and multiplying by total inflorescences for that plant, since we did not always collect all inflorescences. Seed counts were modeled with a negative binomial GLMM, while per-seed mass was modeled with a LMM. Germination data were analyzed as for seed counts, with the number of germinated seeds as the response variable.

For each plant in the 2023 experiment, we calculated the proportion of phenolic offspring and used this as the response variable in a LMM with the same structure as our germination model. Finally, for each plant, we estimated the total number of linalool and phenolic offspring as follows: we first estimated the total seeds produced by multiplying the mean seeds per inflorescence by total number of inflorescences produced. We then multiplied that by the germination rate to calculate the number of offspring and multiplied that by the proportion of linalool and phenolic offspring, respectively, from germination trials. We assessed whether offspring production was influenced by bee infection status and maternal chemotype using a zero-inflated GLMM. Finding a significant maternal chemotype X infection interaction (see *Results*), we then ran separate GLMMs for infected and uninfected tents, asking how predicted offspring number was influenced by maternal chemotype, offspring chemotype, and a maternal X offspring chemotype interaction.

## Acknowledgements

Thanks to the research assistants who contributed to this project, especially Alli Cwalinksi. Jim Cronk supported tent experiments and Baodong Wu supported greenhouse work. Thanks to Koppert Biological Systems for providing discounted research bumble bee colonies, and Ann Arbor Natural Areas Protection and Nasami Farm for plants. This study was funded by the United States National Science Foundation (DBI-2109520 to GF) and the Canadian Natural Science and Engineering Research Council (RGPIN-2024-06190 to GF).

