## Supplementary Material for "Self-medicating behavior in bumble bees has cascading consequences for pollination and plant reproduction"

**S1. SUPPLEMENTAL METHODS**

S1.1. *Additional information about* Monarda fistulosa *biology and source of experimental plants*

Across its range, the most common chemotypes of *Monarda* produce primarily thymol or carvacrol, isomeric monoterpenoid phenolics. Non-phenolic chemotypes, including linalool, geraniol, and 1,8-cineole, are more localized and uncommon (1). Chemotype in the Lamiaceae, presumably including *Monarda*, is under multilocus epistatic control (2). In *Thymus vulgaris* L., the dominance relationship for these chemotypes is: thymol < carvacrol < linalool < geraniol (2); this relationship likely holds true in *Monarda*. *Monarda* inflorescences are tightly packed capitula comprising several dozen to several hundred flowers. Flowers open sequentially, beginning from the center of the capitulum, over the course of 7-14 d, such that at any given time only a fraction of flowers are open.

Most plants used in the study (including all linalool and 1,8-cineole plants) were transplanted in July 2021 or June 2022 as mature plants from wild populations in and around Ann Arbor, MI, USA. A smaller number of plants were transplanted in June 2022 from a single population from Nasami Farm (Whately, MA, USA: 42.461307, -72.642117).

S1.2. *Do dominant monoterpenes from distinct chemotypes differ in their effect on* Crithidia *in vivo?*

S1.2.1 *Preparation of sucrose diets for testing the effect of monoterpenes on* Crithidia *infection*

To make sucrose diets, we followed a three-step process. First, because of the low solubility of several of the monoterpenes, we mixed 1 g of the pure compound with the appropriate volume of 95% ethanol to create a 40% w/w stock solution for each compound. Stock solutions were maintained at -20 ^o^C for the duration of the experiment. Then, once per week during the experiment, we did a two-step dilution of the stock solution with 30% sucrose, first making a 10% v/v monoterpene solution and then drawing 1 μl from this solution to add to 9.999 ml sucrose to create the final 10 ppm solution. Individual diet portions were stored in scintillation vials at -20 ^o^C and thawed at room temperature as needed before transferring to sucrose wells. In addition to the five monoterpene diets, we created a monoterpene-free control by adding a volume of 95% ethanol equivalent to that found in the monoterpene diets to sucrose solution.

S1.2.2*. Assessing sucrose and pollen consumption across monoterpene diets*

To test whether monoterpene diet treatment affected consumption of sucrose solution and/or pollen, and whether consumption affected *Crithidia* infection, we measured sucrose and pollen consumption over the initial 48 h of the trials. At the beginning of each trial, we weighed the sucrose feeder and pollen ball before we introduced each bee to its cup. We repeated these measurements 48 h later when we replaced sucrose and pollen.

To assess the mass of sucrose solution and pollen ball lost to evaporation, we set up evaporation controls (cups identical to those for our experiment, but without a bee), one per diet treatment per date. For each trial, we used data from the evaporation controls to apply a diet-specific evaporation correction to the sucrose and pollen consumption data. To do so, we first regressed final weight (after 48 h) against initial weight for each sucrose diet and for the pollen ball in control cups. We then used the `predict` function to calculate a diet-specific evaporation correction, which we applied to our observed consumption values. We analyzed whether evaporation and evaporation-corrected consumption varied across monoterpene sucrose diet treatments using LMMs, with treatment as fixed effect and date of trial initiation and bee colony as random effects. Where a significant treatment effect was found, we used a post-hoc Tukey’s HSD test to determine differences among the diet treatments.

S1.2.3. *Assessing* Crithidia *infection in bees*

To screen bees for *Crithidia*, we first sacrificed the bees to be screened, then carefully removed the gut contents (including midgut and hindgut). We placed the gut of each bee in an individual 2 ml microcentrifuge tube along with 300 µl of Ringer’s solution and homogenized the gut tissue using a plastic pestle. We let the solution sit for ~4 h to allow *Crithidia* to swim out of the gut tissue, then withdrew 150 μl of the supernatant from each sample into a fresh microcentrifuge tube. After homogenizing this solution with a pipette, we added 10 μl onto a Neubauer hemocytometer and counted the number of active *Crithidia* cells in 0.02 μl using a compound microscope (Olympus Bx42) at 400x.

S1.2.4. *Inoculation of bees with* Crithidia

To ensure that experimental colonies were free of *Crithidia* prior to inoculating experimental bees, we screened 5 worker bees weekly from each experimental colony. To prepare inoculum, we first removed the guts from 5 workers each from two colonies previously infected with *Crithidia* from a strain originally isolated from wild *B. impatiens* from Stone Soup Farm (Hadley, MA, USA: 42.363911, –72.567747) in 2014 and subsequently maintained in commercial colonies of *B. impatiens*. We screened these samples for *Crithidia* using the technique described above (*Assessing* Crithidia *infection in bees*). We selected samples with moderate to high infection intensity (between 25-100 *Crithidia* cells 0.02 μl^-1^) from 1-2 bees from one colony to make inoculum, mixing the remaining 140 μl of supernatant from the chosen sample(s) with additional Ringer’s solution and 50% w/v sucrose solution to create an inoculum with 600 cells µL^-1^ in 25% sucrose. Experimental bees were removed from their colony and starved for 1 h, and each bee was fed a 15 µL droplet of inoculum (9000 *Crithidia* cells). Only bees that consumed the entire droplet were entered into the experiment. Consuming this *Crithidia* cell concentration is within the range of what bees are exposed to in nature from infected feces (3, 4).

S1.3. *Does* Crithidia *infection mediate foraging behavior on* Monarda *chemotypes?*

S1.3.1. *Subcolony preparation for tent experiments*

At least 10 days prior to using a colony, we removed all workers, brood cells, and honey pots from the original colony box and placed half in each of two clean colony boxes (Biobest Group USA, Romulus, MI, USA). One of the boxes was designated ‘uninfected’ and the other ‘infected’. The queen from the original colony was placed in the ‘uninfected’ subcolony. Over the course of the next week, we added *Crithidia* inoculum into partially filled honey pots of the ‘infected’ colony, for a total of 4-8 additions depending on the colony. Inoculum was prepared as described for diet assays. One week after initial inoculation, we dissected 5 workers and assessed *Crithidia* infection as described above. Colony checks continued every two days until at least four of the five workers we checked were infected, at which point we began to use bees from that subcolony in trials.

We retained the queen in the ‘uninfected’ subcolony to extend subcolony longevity (i.e., through continued production of larvae) without compromising the likelihood of bees in the ‘infected’ subcolony being infected via attenuation of infection intensity in later-emerging workers. It is possible that the presence of the queen in ‘uninfected’ but not ‘infected’ subcolonies may have influenced foraging activity by workers. In one study, workers from queenless *Bombus terrestris* colonies showed greater sucrose sensitivity and, as a result, improved olfactory learning, than workers from queenright colonies (5), but it is not known whether the absence of a queen would alter foraging preferences in naïve bees. However, queens in two of the six ‘uninfected’ subcolonies died soon after we established the subcolonies and before we began the experiment (one each in 2022 and 2023), making it unlikely that the differences seen in tents pollinated by bees from ‘infected’ vs. ‘uninfected’ subcolonies are due to the presence or absence of a queen. Moreover, source colony explained relatively little variance in bee foraging behavior in the 2022 experiment, and there was no indication that the bees in the queenless colony showed different preferences for *Monarda* chemotypes or differed in overall foraging activity compared to queenright colonies.

S1.3.2. *Tent construction for experiments*

We constructed foraging tents with 1.3 cm-diameter PVC pipe (Charlotte Pipe, Charlotte, NC, USA) and polyester mosquito netting (Jo-Ann Fabrics, Hudson, OH, USA). Tent frames were either 91cm x 91cm or 61cm x 122cm, with a height of 61cm, 91cm, or 122cm, to accommodate the variation in spacing and height of plants across the site. To allow us to introduce and remove bees from the tents, sleeves made of mosquito netting were affixed to two sides of each tent. Additional mesh fabric at the base of the tents was weighed down to prevent bee escape.

S1.3.3. *Behavioral trials – two-plant experiment*

Each day of trials with paired plants in 2022, we set up trios of worker bees, each comprised of either infected or uninfected bees from a single subcolony. Bees were removed from the subcolony, placed in a mesh-bottomed 16 oz plastic cup, and held at 4^o^C for 15 min to chill. Once bees were chilled, each bee was marked on her thorax with a uniquely colored oil-based paint pen (Sharpie, Atlanta, GA USA). We then placed the trios of bees (hereafter referred to as microcolonies) in a cooler without ice for transport to the field and storage until the trial began. During this time, bees did not have access to food to motivate foraging. In four cases, a bee died before the beginning of the trial; those trials included only two bees. On each day that we conducted observations, bees from at least two parent colonies were used.

Before beginning a trial, we first measured the height of the tallest open inflorescence and counted the number of open inflorescences on each of the two plants selected for observation. We checked the trial arena for flowers other than those on experimental plants and for wild pollinators; when detected, these were removed. We then erected the tent and introduced one trio of bees into the tent, noting the time of introduction. Each trial was conducted by two observers, with each observer focused on one of the two plants. Observers recorded the time of first foraging on their plant for each bee. Once one bee visited both plants, observers began observing all bees, regardless of whether they had visited both plants. If a bee was already foraging at the time of the observation start (i.e., it had begun foraging on one plant but had not yet switched plants), this visit was included, with visit duration and number of probes recorded from the initiation of the observation. If a bee continued to forage after the conclusion of an observation period, the duration and number of probes for that visit were truncated. We initiated observations after one bee had visited both plants so we could be confident that that bee was not choosing to forage on one plant simply because it was not aware of the presence of the other plant. We did not wait for >1 bee to visit both plants because we did not want nectar resources to be depleted from plants prior to the observation. An alternative analysis of foraging behavior including data only from bees that foraged on both plants gave the same results as the analysis including all bees.

Bees generally began foraging immediately upon introduction to the tent. In cases where no bees actively foraged during the first 30 min after introduction, that trial was terminated, and those bees were immediately removed from the tent. In cases where bees were actively foraging but no bee had switched between plants during the first 30 min, we began the observation period at the 30 min mark, but noted that no bee had, at that point, visited both plants.

Seven bees that we inoculated with *Crithidia* did not show evidence of infection when dissected. In our analysis of foraging behavior, these bees were treated as uninfected. Since we never had >1 *Crithidia*-free bee per group of infected bees, we treated those tents as infected for our analyses of stigma pollen load.

S1.4. *Does infection-mediated foraging behavior differentially affect pollination and plant female reproduction across chemotypes?*

S1.4.1. *Assessing pollen deposition – two-plant experiment (2022)*

Prior to the onset of flowering, we randomly selected inflorescences for assessing pollination, marked these with a piece of white labeling tape (Thermo Fisher Scientific, Waltham, MA, USA) affixed to the stem directly below the bracts, and enclosed them in a 13 cm x 17 cm mesh drawstring bag (Uline, Pleasant Prairie, WI, USA). For inflorescences assigned to a given observation, we removed mesh bags after the foraging tent was erected but before the bees were introduced, and we wrote the date of observation on the tape tag. After we removed bees from the tent at the end of the trial, we collected one receptive stigma from each unbagged inflorescence to assess pollen deposition. We identified receptive stigmas by the presence of bifurcated stigma lobes (6); from those flowers with receptive stigmas, one was chosen at random and clipped at the base of the stigma. The number of pollen grains on the stigma was quantified as described in the main manuscript.

S1.4.2. *Seed production – two-plant tent experiment (2022)*

We harvested inflorescences as they matured, between 11 Aug and 01 Sep 2022. Upon harvest, bagged inflorescences were left in the mesh bag to contain any dislodged seeds. We subsequently measured the diameter of the mature inflorescence, counted the number of seeds, weighed the group of seeds, and divided this by the number of seeds to calculate per-seed mass.

We assessed female reproductive success as described for the 6-plant trials (see *Methods*, main manuscript), with the following modifications: 1) We did not assess seeds produced per plant. 2) The number of receptive female flowers at the time of the pollination trial was included as a fixed effect in all models. 3) Model random effects included only plant ID nested within plot. We had insufficient data to compare seed measures in phenolic-1,8-cineole tents (N = 16 1,8-cineole inflorescences).

S1.5. *Comparison of floral traits across chemotypes*

To determine whether floral traits differed between chemotypes, we assessed the following floral traits from experimental plants at our field site: inflorescence number, inflorescence width, flower number per inflorescence, and corolla length, width and height. Inflorescence number was measured from experimental plants in the 2023 trials (in 2022, plants began the season at very different sizes, biasing measures of inflorescence number); all other traits were assessed in both 2022 and 2023 from plants that were used in the 2022 trials and remained on site but were not used in 2023.

We measured floral traits from three flowers from 1-5 inflorescences per plant, from 32 different plants (8 plants per chemotype); the total number of flowers measured varied depending on availability at the time traits were measured (carvacrol: N = 60; linalool: N = 30; thymol: N = 30; 1,8-cineole: N = 30). With digital calipers, the inflorescence diameter of each inflorescence was measured to the nearest 0.01 mm. We then chose three flowers per inflorescence at random and measured their corolla length, corolla width, and floral diameter to the nearest 0.01 mm.

To test whether floral traits differed across chemotypes, we used linear mixed-effects models (corolla measurements, inflorescence width) and negative-binomial generalized linear mixed-effects models (inflorescence number), with chemotype and sampling year as fixed effects and plant ID as a random effect. We checked conformity to assumptions with DHARMa.

S1.6. *Assessing nectar monoterpene concentration*

In July 2023, we collected a single sample of nectar from each chemotype (thymol and linalool). To do so, we first enclosed plants of a known chemotype under a mesh tent to exclude nectar consumers for at least 24h before sampling. To extract nectar, we inserted a 25 μl microcapillary tube (Drummond Scientific, Broomall, PA, USA) into the corolla mouth and down to the base of the flower, where nectaries are located. Because we needed approx. 20 μl of nectar per sample, and *Monarda* flowers produce very little nectar per flower, we had to probe many flowers, from multiple plants, to extract the necessary volume. We calculated the volume of nectar extracted by measuring the height of the sample in the tube and using the formula $v=\pi r^{2}h$, where $r$ is the interior radius of the microcapillary tube and *h* is the height in mm of the sample in the tube. The sample was then expelled into a 1.5 ml microcentrifuge tube containing 80 μl of 100% ethanol. Upon returning to the lab, samples were stored at -20°C until gas chromatography-mass spectrometry (GC-MS) analysis was performed.

The GC-MS was performed with a single quad 5977C MS (Agilent Technologies) linked to an Agilent 8890 GC (Agilent Technologies). The GC-MS was equipped with a DB5 capillary column (30 m × 0.25 mm ID; J&W, Agilent Technologies). Before injection, samples were concentrated to 50 μl, from which aliquots of a 4 μl sample were injected in split mode (1:1) at an oven temperature of 50°C. After 1 min, the splitter valve was opened and the temperature was increased at a rate of 10°C min^−1^ to 310°C. A constant flow of 2 ml min^−1^ helium was used as the carrier gas. For quantitative analysis, 1 ng of carvacrol was used as an external standard. Identifications of carvacrol and thymol were performed by comparing the mass spectra of natural samples with those of commercial databases (NIST library, ADAMS) and with the spectra of synthetic compounds. Absolute concentration of the terpenes was conducted based on the signal intensity of the external standards.

S1.7. *Testing pollinator dependence*

S1.7.1. *Data collection*

In 2022, prior to the onset of flowering, we randomly selected inflorescences for inclusion in pollination trials, and assigned chosen inflorescences to one of three treatments: bagged (covered with a mesh bag for the entirety of the bloom period; received no pollination), tent-pollinated (bagged except for a single trial, in which the mesh bag was removed to allow visitation by experimental bees of known infection status; these are the same inflorescences used to assess the role of chemotype and infection status on pollination, see main text), or open-pollinated (unbagged for the duration of the bloom period; open to all pollinators). Inflorescences were randomly assigned to treatment within a plant; each plant included at least one inflorescence assigned to each treatment. The number of inflorescences included in each treatment differed among plants in proportion to the total number of inflorescences each plant produced. All inflorescences were marked with a piece of tape affixed to the stem just below the bracts. We assessed pollen deposition (see *Assessing pollen deposition* above) on a single stigma from each inflorescence in the bagged and open-pollinated treatments.

S1.7.2. *Data analysis*

Because stigma pollen data were strongly zero-inflated, we used hurdle models as described in the main manuscript to analyze whether stigma pollen load was influenced by pollination treatment [bagged, open only during foraging trials (“tent-pollinated”), and open to all visitors (“open-pollinated”)]. To test for differences among treatments, our model included *Monarda* pollen deposition as the response variable, inflorescence treatment and chemotype as fixed effects, and date of stigma collection and plot as random effects.

S2. **SUPPLEMENTAL RESULTS**

S2.1. *Seed production – two-plant tent experiment (2022)*

Comparing phenolic and linalool tent-pollinated inflorescences, there was no effect of chemotype, infection status, or chemotype × infection status interaction on either seed number or mass per seed after controlling for the effects of inflorescence size, inflorescence number, and plant size (Table S1). The number of receptive flowers open during the foraging trial did not influence seed number (ꭓ^2^ = 0.18, df = 1,89, p = 0.7), and this variable was not included in the final model. This indicates that pollination by experimental bees did not contribute substantially to seed production, an inference that is further supported by the lack of difference in seed production between tent-pollinated and bagged inflorescences. As such, the seed production results likely do not reflect the impact of differential foraging preference due to *Crithidia* infection. Therefore, we conducted the six-plant tent experiments to allow for more realistic levels of pollination and better assess effects of infection-mediated foraging preferences on plant reproduction.

S2.2. *Results of pollinator dependence tests*

We assessed stigma pollen deposition from 12 bagged flowers and 19 open-pollinated flowers, and compared these with stigma pollen deposition from the 159 tent-pollinated flowers we collected. Inflorescence treatment did not significantly influence the likelihood of a stigma receiving at least one *Monarda* pollen grain (Table S1; Fig. S2A), but for stigmas that received *Monarda* pollen, treatment significantly influenced pollen receipt (Table S2). Bagged stigmas had significantly more pollen grains than either open- or tent-pollinated stigmas (p = 0.009 for both contrasts), while open- and tent-pollinated stigmas did not differ in pollen receipt (p = 0.7; Figure S2B), indicating substantial autogamous pollination. For bagged flowers, there was no effect of chemotype on stigma pollen receipt (ꭓ^2^ = 1.9, d.f. = 2, 7, p = 0.4). For open-pollinated flowers (i.e., those accessible to all pollinators throughout the bloom period), stigma pollen load differed among chemotypes (ꭓ^2^ = 6.8, d.f. = 2, 18, p = 0.03), with thymol plants receiving significantly more pollen than linalool plants (2.2±1.2 vs. 0.5±0.4 pollen grains on stigma, p = 0.04), and carvacrol plants intermediate (3.6±3.3 pollen grains on stigma) and not significantly different from either thymol or linalool plants.

We assessed seed production and per-seed mass from 52 bagged inflorescences and 57 open-pollinated inflorescences, and compared these with data from the 150 tent-pollinated inflorescences. Inflorescence treatment significantly impacted the number of seeds produced per inflorescence, with open-pollinated inflorescences producing >3-fold times more seeds than bagged or tent-pollinated inflorescences (Table S1, Figure S3C). This was despite bagged inflorescences having higher stigma pollen loads than open-pollinated ones, indicating that *Monarda*, while self-fertile, benefits strongly from cross-pollination. There was no difference in seeds per inflorescence between bagged and tent-pollinated inflorescences (Figure S3C), indicating that the contribution of foraging trials to pollination was minimal. Per-seed mass did not differ among treatments (Table S1).

S2.3. *Germination trials – six-plant experiment (2023)*

Germination rates were low ($\bar{x}$ = 0.32±0.19 seedlings seed^-1^) and positively correlated with maternal plant height, but not other predictors (Table 3, Fig. 4C). Offspring chemotype was strongly biased towards phenolic (i.e., medicinal) plants (71.0% of offspring), and this was influenced by maternal chemotype (50.3% of offspring were phenolic for linalool plants, vs. 87.5% for thymol plants). Neither infection status nor chemotype $\times$ infection status significantly influenced the proportion of phenolic offspring from germinated seeds (Table 3; Fig. 4D).

**SUPPLEMENTARY TABLES**

Table S1. *Monarda* stigma pollen deposition, seed production, and per-seed mass from pollinator dependence trials and two-plant tent experiments.

| **Response**  Predictor | **ꭓ^2^** | **DF** | **p** |
| --- | --- | --- | --- |
| **Likelihood of having ≥1 pollen grain on stigma, pollinator dependence experiment** | | | |
| Pollination treatment^1^ | 4.24 | 2, 205 | 0.1 |
| **Number of pollen grains on stigma, pollinator dependence experiment** | | | |
| *Pollination treatment*^1^ | *9.11* | *2, 86* | *0.01** |
| **Seeds per inflorescence, pollinator dependence experiment** | | | |
| *Pollination treatment*^1^ | *58.12* | *2, 197* | *<0.001**** |
| *Inflorescence diameter* | *54.42* | *1, 197* | *<0.001**** |
| *Observer identity* | *10.07* | *3, 197* | *0.02** |
| **Per-seed mass, pollinator dependence experiment** | | | |
| Pollination treatment^1^ | 4.00 | 2, 186 | 0.5 |
| *Chemotype*^2^ | *8.12* | *2, 186* | *0.02** |
| **Seeds per inflorescence, two-plant tent experiment: phenolics vs. linalool** | | | |
| Chemotype | 1.53 | 1, 97 | 0.2 |
| Infection status | 0.40 | 1, 97 | 0.5 |
| Chemotype × infection status | 0.00 | 1, 97 | >0.9 |
| *Inflorescence diameter* | *16.56* | *1, 97* | *<0.001**** |
| *Total # of inflorescences on plant* | *10.80* | *1, 97* | *0.001*** |
| *Plant size* | *4.07* | *1, 97* | *0.04** |
| **Per-seed mass, two-plant tent experiment: phenolics vs. linalool** | | | |
| Chemotype | 3.58 | 1, 90 | 0.06• |
| Infection status | 1.20 | 1, 90 | 0.3 |
| Chemotype × infection status | 0.50 | 1, 90 | 0.5 |
| *Total # of inflorescences on plant* | *7.38* | *1, 90* | *0.007*** |

Italics indicate a significant effect (p < 0.05); •p<0.1, *p<0.05, **p<0.005, ***p<0.001.

^1^Treatments for pollinator dependence experiment included bagged (all pollinators excluded throughout flowering), tent-pollinated (inflorescence available for pollination only during a single tent foraging trial, and open-pollinated (inflorescence always available; ambient pollination). ^2^Significant effect of chemotype due to per-seed mass for 1,8-cineole plants being higher than other chemotypes, which were indistinguishable from one another.

Table S2. Floral traits did not differ among *Monarda* chemotypes. *P-*values are not corrected for multiple testing.

| Trait | Trait value (mean±s.e.) | | | | Test statistic | df | *P* |
| --- | --- | --- | --- | --- | --- | --- | --- |
|  | Carvacrol | 1,8-Cineole | Linalool | Thymol |  |  |  |
| Inflorescence number | – | – | 35.7±4.6 | 38.2±4.1 | 0.41^1^ | 1, 23 | 0.7 |
| Inflorescence width (mm) | 48.15±1.71 | 50.46±3.21 | 45.41±1.41 | 51.73±1.81 | 1.72^1^ | 3, 50 | 0.3 |
| Corolla length (mm) | 17.72±0.37 | 18.65±0.36 | 17.11±0.34 | 16.82±0.36 | 6.8^2^ | 3,158 | 0.08 |
| Corolla width (mm) | 2.02±0.11 | 1.92±0.06 | 2.00±0.08 | 1.65±0.05 | 6.2^2^ | 3,158 | 0.1 |
| Corolla flare (mm) | 11.81±0.33 | 12.32±0.50 | 14.56±0.61 | 11.71±0.32 | 5.3^2^ | 3,158 | 0.2 |

^1^z-score. ^2^ $\chi$^2^.

Table S3. Catalog numbers for chemicals used in assay of effect of monoterpenes on *Crithidia* infection in *Bombus impatiens* workers.

| **Compound** | **Sigma Aldrich catalog number** |
| --- | --- |
| Carvacrol | 282197 |
| Geraniol | 48798 |
| Linalool | 51782 |
| Thymol | T0501 |
| 1,8-Cineole | 29210 |

Table S4. Full formulations for statistical models used in data analysis. Boldface indicates a fixed effect that was retained in the best model.

| **Model** | **Dependent variable** | **Fixed effects** | **Random effects** | **Error distribution** |
| --- | --- | --- | --- | --- |
| Effect of monoterpenes on *Crithidia* infection | *Crithidia* cell counts | **Diet**  Sucrose consumption  (evaporation-corrected)  Pollen consumption  (evaporation-corrected)  Marginal cell length | Colony of origin  Inoculation date | Negative binomial |
| Effect of chemotype and infection on foraging (hurdle model) | Bee visitation (binary Y/N) | **Plant chemotype**  **Bee infection status**  **Chemotype × infection status**  Marginal cell length  Plant height^1^  Inflorescence number | Plot / Plot-Date^2^  Colony of origin | Binomial |
| Effect of chemotype and infection on foraging (hurdle model) | Bee visit duration in seconds | **Plant chemotype**  **Bee infection status**  **Chemotype × infection status**  Marginal cell length  Plant height  Inflorescence number | Plot / Plot-Date^2^  Colony of origin | Zero-truncated negative binomial |
| Effect of chemotype and infection on pollination (hurdle model) | Presence of pollen on stigma (Y/N) | Plant chemotype  Bee infection status  Chemotype × infection status  Marginal cell length  Plant height  # of inflorescences with open flowers | Plot / Plot-Date^2^  Colony of origin | Binomial |
| Effect of chemotype and infection on pollination (hurdle model) | *Monarda* pollen grain count for stigmas with pollen | Plant chemotype  Bee infection status  Chemotype × infection status  Marginal cell length  Plant height  # of inflorescences with open flowers | Plot / Plot-Date^2^  Colony of origin | Zero-truncated negative binomial |
| Effect of chemotype and infection on seed production | Seeds per inflorescence | **Maternal chemotype**  **Bee infection status**  **Chemotype × infection status**  **Inflorescence diameter**  Plant height  **Observer** | Tent / Plant ID  Clone source ID^3^ | Negative binomial |
| Effect of chemotype and infection on seed production | Seeds per plant | **Maternal chemotype**  **Bee infection status**  **Chemotype × infection status**  **Mean inflorescence diameter**  Plant height | Tent / Plant ID  Clone source ID^3^ | Negative binomial |
| Effect of chemotype and infection on seed mass | log_10_(Per-seed mass) | **Maternal chemotype**  **Bee infection status**  **Chemotype × infection status**  **Mean inflorescence diameter**  Plant height  **Observer** | Tent / Plant ID  Clone source ID^3^ | Gaussian |
| Effect of chemotype and infection on germination rate | Proportion of seeds germinated | **Maternal chemotype**  **Bee infection status**  **Chemotype × infection status**  Inflorescence diameter  **Plant height** | Tent  Clone source ID^3^ | Gaussian |
| Effect of chemotype and infection on offspring chemotype | Proportion of offspring that are phenolic | **Maternal chemotype**  **Bee infection status**  **Chemotype × infection status**  Inflorescence diameter  Plant height | Tent  Clone source ID^3^ | Gaussian |
| Effect of chemotype and infection on offspring number | Estimated number of offspring, separated by chemotype | **Maternal chemotype**  **Bee infection status**  **Maternal chemotype × infection status**  Offspring chemotype  Maternal chemotype × Offspring chemotype  **Inflorescence diameter**  **Plant height** | Clone source ID^3^ / Maternal plant ID | Zero-inflated negative binomial^4^ |
| Effect of chemotype and infection on phenolic offspring number | Estimated number of phenolic offspring | **Maternal chemotype**  **Bee infection status**  **Chemotype × infection status**  **Inflorescence diameter**  **Plant height** | Clone source ID^3^ | Negative binomial |
| Effect of chemotype and infection on linalool offspring number | Estimated number of linalool offspring | **Maternal chemotype**  **Bee infection status**  **Chemotype × infection status**  **Inflorescence diameter**  **Plant height** | Clone source ID^3^ | Negative binomial |

^1^Included in final model for comparison between phenolic and 1,8-cineole plants but not between phenolic and linalool plants (see Table 1, main manuscript).

^2^Dummy variable to account for paired structure of two plants per tent.

^3^ID of the plant from which the experimental plant was split, to reflect paired design of plants between paired tents.

^4^Model included an intercept-only zero-inflation component.

Table S6. York University Greenhouse soil recipe.

| Component | Quantity |
| --- | --- |
| Potting Soil (Berger BM1) | 107 L |
| Garden Soil (Alltreat Farms 3-Way Mix) | 15 kg |
| Cattle manure (Alltreat Farms) | 9 kg |
| Sand (Sakrete Play Sand) | 13 kg |
| Water | 7 L |

Table S6. Interactive effects of chemotype and infection status on bumble bee foraging behavior and pollination in two-plant trials in 2022, separately considering thymol and carvacrol plants. Small sample size precluded comparison of pollen deposition between carvacrol and linalool.

| **Predictor** | **z-score** | **df** | ***P*** |
| --- | --- | --- | --- |
| **Visitation likelihood, thymol vs. linalool** | | | |
| Chemotype | 0.85 | 1, 152 | 0.4 |
| Infection status | 0.18 | 1, 152 | 0.7 |
| Chemotype × infection status | 1.79 | 1, 152 | 0.18 |
| *Plant height* | *8.99* | *1, 152* | *0.003*** |
| **Foraging duration, thymol vs. linalool** | | | |
| Chemotype | 0.47 | 1, 119 | 0.5 |
| Infection status | 1.77 | 1, 119 | 0.18 |
| Chemotype × infection status | 0.04 | 1, 119 | 0.8 |
| Plant height† | 0.02 | 1, 119 | 0.8 |
| **Visitation likelihood, carvacrol vs. linalool** | | | |
| *Chemotype* | *4.35* | *1, 126* | *0.04** |
| Infection status | 0.09 | 1, 126 | 0.8 |
| Chemotype × infection status | 0.66 | 1, 126 | 0.4 |
| *Plant height* | *4.56* | *1, 126* | *0.03*** |
| **Foraging duration, carvacrol vs. linalool** | | | |
| *Chemotype* | *5.65* | *1, 106* | *0.02** |
| Infection status | 0.48 | 1, 106 | 0.5 |
| Chemotype × infection status | 0.66 | 1, 106 | 0.4 |
| **Likelihood of having ≥1 pollen grain, thymol vs. linalool** | | | |
| Chemotype | 0.05 | 1, 75 | 0.8 |
| Infection status | 0.15 | 1, 75 | 0.7 |
| Chemotype × infection status | 0.10 | 1, 75 | 0.8 |
| **Number of *Monarda* pollen grains on stigma, thymol vs. linalool** | | | |
| Chemotype | 0.33 | 1, 21 | 0.6 |
| Infection status | 0.47 | 1, 21 | 0.5 |
| *Chemotype × infection status* | *4.98* | *1, 21* | *0.03** |

Italics indicate a significant effect (p < 0.05); *p<0.05, **p<0.005.

Table S7. Interactive effects of chemotype and *Crithidia* infection on bumble bee foraging behavior in two-plant experiment in 2022: number of inflorescence probes and number of plant visits.

| **Predictor** | **z-score** | **df** | ***P*** |
| --- | --- | --- | --- |
| **Number of inflorescence probes, phenolic vs. linalool^1^** | | | |
| Chemotype | 2.22 | 1,266 | 0.1 |
| Infection status | 0.09 | 1,266 | 0.7 |
| Chemotype × infection status | 1.21 | 1,266 | 0.3 |
| **Number of plant visits, phenolic vs. linalool^1^** | | | |
| Chemotype | 2.08 | 1,266 | 0.1 |
| Infection status | 1.68 | 1,266 | 0.2 |
| Chemotype × infection status | 0.69 | 1,266 | 0.4 |
| **Number of inflorescence probes, phenolic vs. 1,8-cineole^1^** | | | |
| Chemotype | 0.84 | 1,64 | 0.4 |
| Infection status | 0.77 | 1,64 | 0.4 |
| Chemotype × infection status | 0.38 | 1,64 | 0.5 |
| **Number of plant visits, phenolic vs. 1,8-cineole^1^** | | | |
| Chemotype | *0.28* | 1,64 | 0.6 |
| Infection status | 1.26 | 1,64 | 0.3 |
| Chemotype × infection status | 0.25 | 1,64 | 0.6 |

^1^ These models assess data only from plants that were visited at least once, and therefore use zero-truncated negative binomial error distributions. The models for probability of visitation are identical to that seen in Table 1.

Figure S1. Effect of monoterpene sucrose diets on *Crithidia* infection *in vivo*. Large points indicate estimated marginal mean; bars represent ±1SE. Small points represent individual bees. Letters indicate statistical differences from post-hoc analysis.

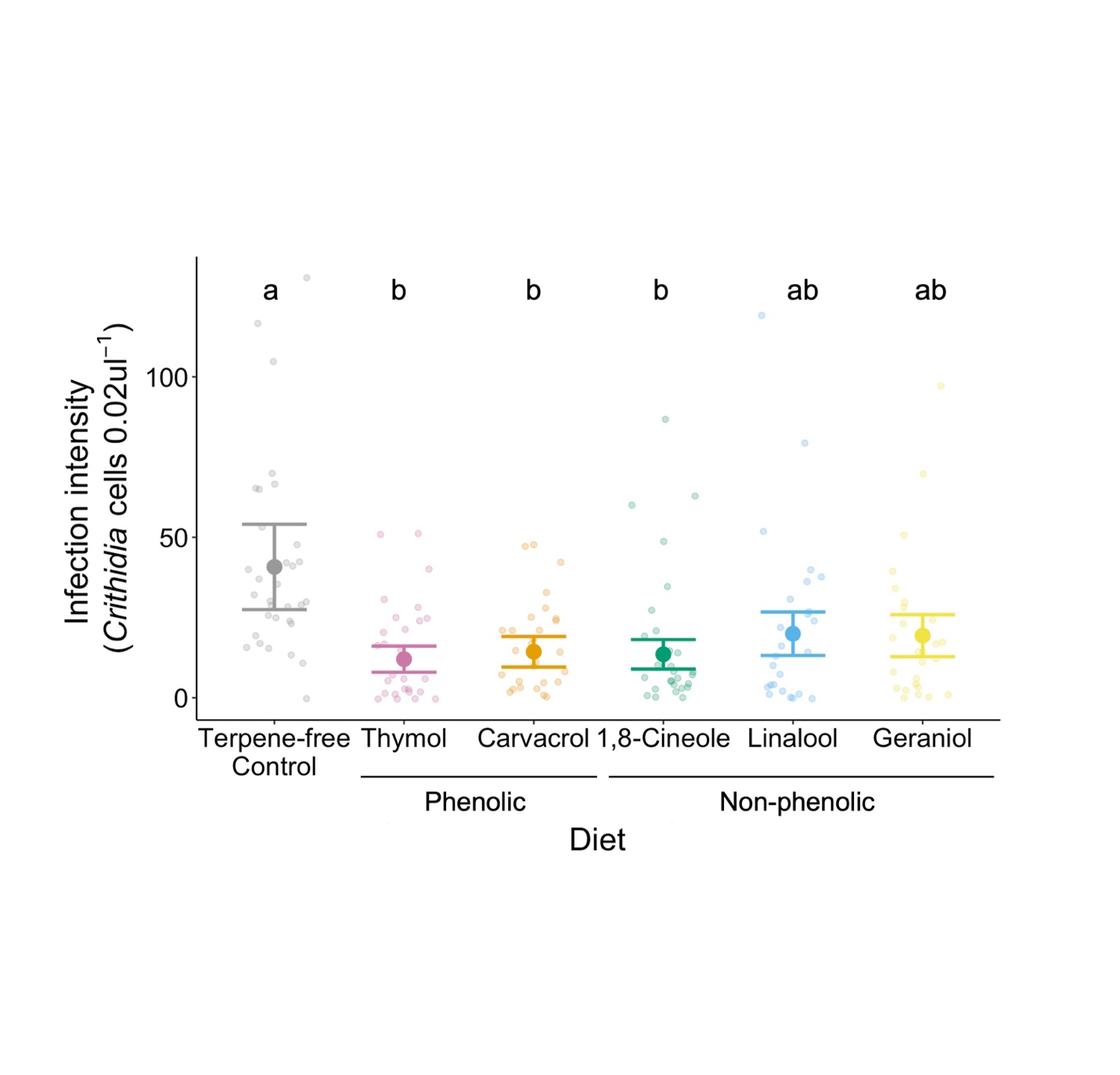

Figure S2. Influence of pollination treatment on A) likelihood of a stigma receiving pollen, B) stigma pollen deposition (number of pollen grains per stigma), and C) seeds produced per inflorescence in two-plant trials in 2022. Pollination treatments: Bagged = all pollinators excluded, Open-pollinated = accessible to all pollinators, Tent-pollinated = open to pollinators only during experimental trials. Letters indicate significant differences among treatments; points indicate means, bars indicate standard error, whiskers indicate 95% confidence interval.

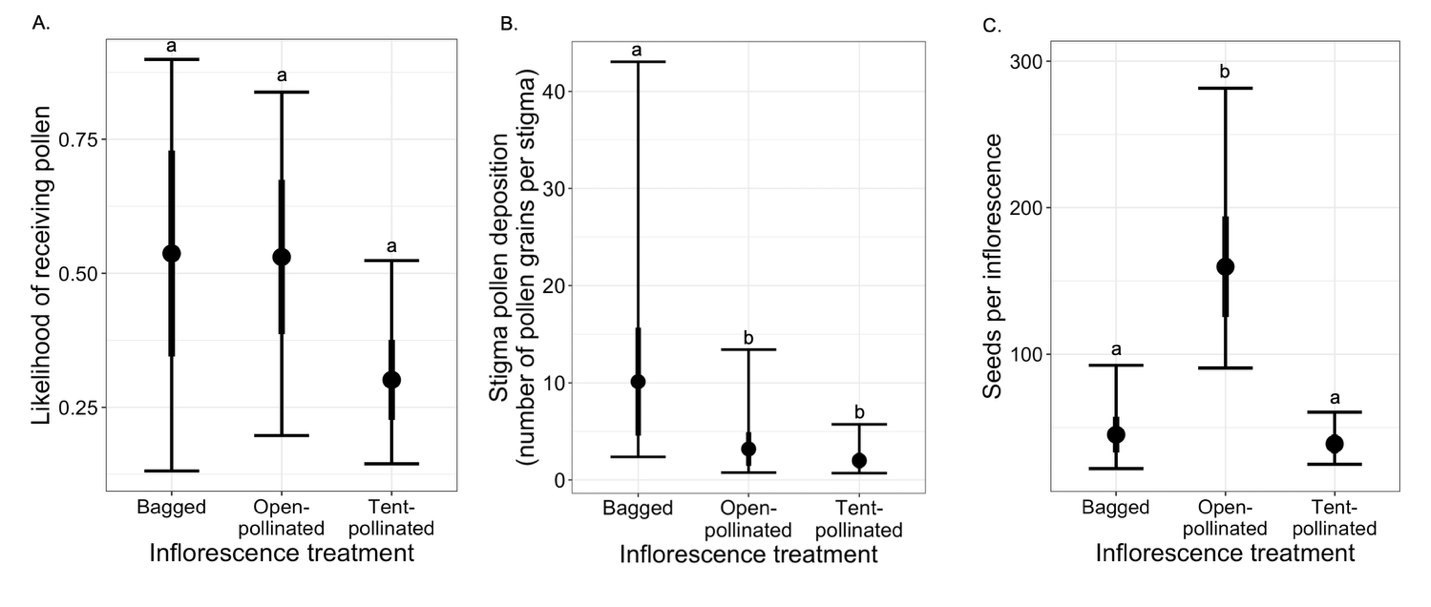
